# Benchmarking long-read genome assemblers for three sequencing protocols and three agricultural species

**DOI:** 10.1101/2025.02.14.638238

**Authors:** Clément Birbes, Andreea Dréau, Denis Milan, Carole Iampietro, Christine Gaspin, Cécile Donnadieu, Matthias Zytnicki, Christophe Klopp

## Abstract

Today, long read technologies make it possible to produce telomere-to-telomere genome assemblies. These assemblies permit more accurate whole genome analyses, including on repeated regions.

The genome assembly process can chain several steps such as read correction, contigs assembly, contig polishing, scaffolding and gap filling, among others. Each step usually requires at least one software package, and input sequences. For the end user, it is not necessarily clear which combination of tools, together with read types, will work best for a given genome.

In this work, we will focus on contig production, which is a central and complex task of the assembly process. It aims at producing the longest, errorless, sequences, called *contigs*, from reads. While it is possible to produce contigs from short reads, long reads are now widely preferred, since they produce much longer contigs.

In this work, we evaluate several contig producing software packages (usually named *assemblers*), on long reads generated by two sequencers using three protocols, for three eukaryotic, complex species with different characteristics. Our aim is twofold. First, we would like to give readers insight on the impact of sequencing technology and assembler combinations, in order to help them make their choice for a given genome. Second, we would like to present different assembly metrics and provide a critical view on their interpretation.

## Introduction

Today many studied species benefit from at least one reference genome assembly. A genome assembly is a text file usually in FASTA format containing sequences representing chromosomes or chromosome parts. Genome assemblies enable a large set of genomic analysis, including gene expression quantification, genomic variation calling, ChIP-seq peak discovery, methylation state measurement, etc.

With the advent of long read sequencing technologies provided by companies such as PacBio and Oxford Nanopore Technologies, the assembly of small genomes up to tens of megabases is a solved question and the assembly of large genome has been tremendously simplified. It can now be performed by a single researcher for a few thousand dollars compared to the large consortia needed twenty or thirty years ago. It is thus now possible to study new levels of genomic complexities, at a reasonable price.

Genomes can differ in several ways, some of which can be linked. Size has been mentioned before and repeat content is often linked to size. The longer the repeats and the more resembling the more difficult is their disentanglement, and therefore the assembly. Heterozygocity is another characteristic impacting genome assembly. The lower the heterozygocity the easier the assembly, explaining why some projects use double haploids. Recent whole genome duplication will also render assembly more difficult because only few variations are found between chromosomes sharing the same ancestor. These cases being rare, they are often not taken into account in the assembly software packages.

The aim of genome assembly is to produce a *bona fide* representation of the genome. Many strategies can be used, but most now rely on long reads. These reads, which can contain errors at varying degrees, can first be corrected using other reads or on themselves. They are then assembled together, in order to form longer sequences. These sequences can then be polished in order to remove errors and scaffolded into longer pseudo-molecules. In this article, we will only focus on the contig assembly part.

In principle, an assembler takes a set of input reads produced by a sequencer, and stitches them using common sequence part or sequence overlaps, in order to produce the longest possible contigs. Long read assemblers rely on two different methods: Overlap Layout Consensus (OLC) graphs, or de Bruijn graphs. OLC assemblers are comparing the reads to themselves in order to build an overlap graph. De Bruijn assemblers also use a graph, which is built from k-mers, which are overlapping read chunks. In principle, each path should represent chromosome parts called *contigs*, which are extracted from the graph. Although simple at first glance, genome assembly is a highly complex task. Repetition and sequencing errors tend to add spurious information in often very large graphs, which become highly tangled, and detecting this spurious information is far from straight-forward. Moreover, contigs should be as long as possible, and contain as few errors as possible. Likewise, the tools should be fast, and not consume too much memory. As a consequence, each tool implements different strategies in order to solve this multi-objective problem.

Given the wealth of tools, it is necessary to assess their performances through the quality of the produced assemblies. Here again, several metrics can be used. Some metrics only measure the contig sizes. Other metrics compare the assembly with a reference. This reference can be an estimated genome size, a reference genome, a set of reference genes, or the reads themselves. All these references give a fraction of information, which can be used to assess assembly quality. Taken individually, these metrics can be misleading. Our aim is to describe and show their limitations, and to suggest a set of metrics that can faithfully assess the assembly quality.

This article is based on long read datasets produced during the SeqOccIn project which was focused on species of agronomic interest. This project aimed at comparing long read sequencing technologies to build state of the art pipelines for several applications, including genome assembly, variation and methylation calling as well as meta-genomic analysis. This article will include read sets from three species : quail (*Coturnix Japonica*), maize (*Zea mays*) and cattle (*Bos taurus*) and from three sequencing protocols Oxford Nanopore technologies R9.4, PacBio CLR and PacBio HiFi. We present the result of 39 assemblies performed using eight software packages (flye, hifiasm, ipa, lja, NextDenovo, raven, shasta, wtdbg2).

The species have been chosen because of their agronomic interest and their diversity. They have different genome sizes and repeat content. One genome harbors micro-chromosomes which are known to be difficult to sequence and assemble. They all have a reference genome assembly available at the NCBI and EMBL/EBI.

## Material and methods

### Estimated genome sizes

The three genomes vary in size. Although reference assemblies are available, we wanted to use an external, more reliable, estimation of genome size. We used estimations based on flow cytometry, ultraviolet microscopy or biochemical analysis. Figures provided by the Animal Genome Database (Gregory, 2023) and MaizeGDB (Bennett, 1995) are:

- *Bos taurus*: 3.08–3.84 Gbp
- *Coturnix japonica*: 1.26–1.38 Gbp
- *Zea mays*: 2.41–2.74 Gbp

We chose the average values, which are, respectively, 3.53 Gbp, 1.32 Gbp, and 2.57 Gbp.

### Overview of current reference assemblies

The genomes have specific characteristics presented in Table 1.

**Table 1.** Metrics on the current assemblies, computed by the NCBI.

|  | *Bos taurus* | *Coturnix japonica* | *Zea mays* |
| --- | --- | --- | --- |
| Assembly name | ARS-UCD1.2 | Coturnix japonica 2.1 | Zm-B73-REFERENCE-NAM-5.0 |
| Assembly size | 2,715,853,792 | 927,656,957 | 2,182,786,008 |
| # scaffolds | 2,211 | 2,531 | 687 |
| Scaffold N50 | 103,308,737 | 2,975,000 | 226,353,449 |
| # chromosomes and plasmids | 31 | 32 | 12 |
| # protein-coding genes | 21,039 | 16,026 | 34,337 |
| # non protein-coding genes | 9,357 | 4,736 | 10,366 |
| # mRNAs | 63,675 | 40,881 | 57,304 |
| % repetition | 49.66% | 7.82% | 78.51% |
| reference | (NCBI, 2023a) | (NCBI, 2023b) | (NCBI, 2023c) |

All three genomes have already been assembled and released several times. The NCBI provides several statistics on these assemblies, some of which are provided in Table 1. These assemblies have all been generated using long read technologies. A first observation is that the assembly sizes are smaller than the expected genome size, which is commonly observed. Second, none of them are assembled telomere-to-telomere, and the numbers of scaffolds range from 687 to 2,531. Third, the percentage of repetition greatly varies between organisms from 8% to 79%. Although we expect highly repeated genomes to be more difficult to assemble, the variability in repetition is crucial: it is much more difficult to assemble genomes that contain exact repetitions than genomes where repetitions have accumulated errors during evolution.

### Assembly quality metrics

Several metrics can be used to assess the quality of an assembly. They include the NG50 and the LG50, which assess assembly contiguity: the NG50 will be high if the contigs are very long. However, a high NG50 does not imply a good assembly. An assembler that would stitch together contigs from different *loci* would also produce a high NG50, at the expense of the assembly quality.

Other approaches compare the assembly with a reference that is deemed reliable. We first computed the relative size, which is 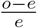, where *o* is the observed size of the assembly, and *e* is the expected size, obtained from cytometry. The relative size is positive if the assembly is longer than the expected one, and negative otherwise.

We also computed coverage metrics, provided by Inspector (Y Chen, Zhang, et al., 2021). It uses a reference assembly, maps the contigs to this assembly, and provided the proportion of the genome which is covered. This metrics is meaningful if the assembly and the reference genome are close enough.

The Merqury package (Rhie, Walenz, et al., 2020) provides several metrics that compare the assembly with the reads themselves. The QV is an estimation of the probability of error for the assembly at the nucleotide level, computed as a Phred score. The completeness estimates the quantity of the nucleotides that have been used in the assembly, divided by the number of nucleotides that have been sequenced. Inspector (Y Chen, Zhang, et al., 2021) also computes the number of assembly errors, categorized as expansions (gaps in read alignment), collapses (extra sequence in reads), and inversions (inverted read alignments). In our analysis, we grouped them as errors. Of note, these metrics based on reads are biased towards the sequencing quality. If the sequencer produces (non-random) errors, these metrics can be high, even though the assembly is far from the actual genome.

The BUSCO tool (Feron and Waterhouse, 2022) uses a list of genes that have been manually curated, and are expected to be present in exactly one copy in the phylum of the species. The *complete*, or BUSCO C, score is the percentage of genes that are found once in the assembly, without major error. The *duplicate*, or BUSCO D, score is the percentage of genes that are found more than once in the assembly. These BUSCO scores vary greatly with the base level quality of the assembly: a few nucleotide errors in an assembled gene may truncate the open reading frame, and hinder BUSCO to detect this gene.

Table 2 links metrics with the types of misassemblies it can detect.

**Table 2.**
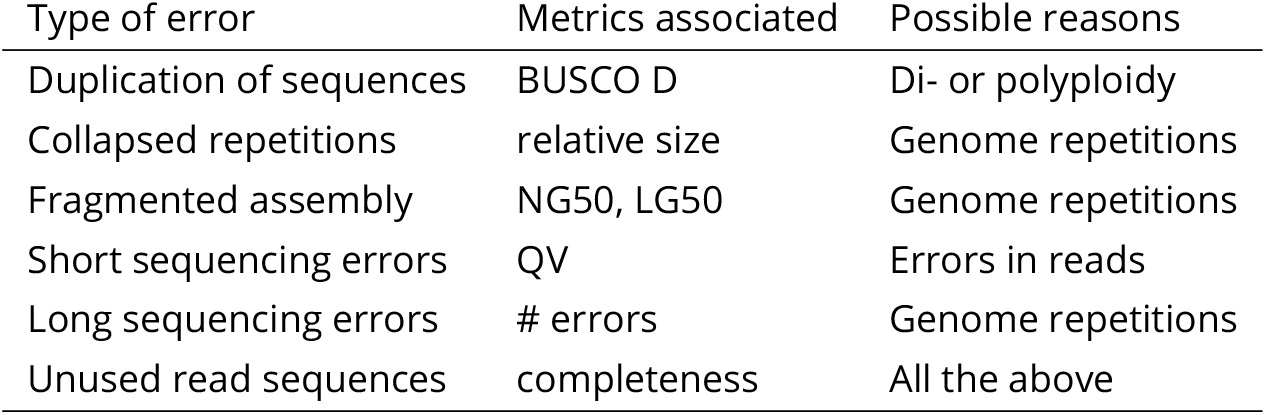
Possible sources of errors in genome assemblies.

| Type of error | Metrics associated | Possible reasons |
| --- | --- | --- |
| Duplication of sequences | BUSCO D | Di- or polyploidy |
| Collapsed repetitions | relative size | Genome repetitions |
| Fragmented assembly | NG50, LG50 | Genome repetitions |
| Short sequencing errors | QV | Errors in reads |
| Long sequencing errors | # errors | Genome repetitions |
| Unused read sequences | completeness | All the above |

The time spent, as well as the memory used by each tool, has also been added to the list of metrics. They can be used to choose whether a given tool can be used, or not, on the available computers. In order to compare the results on the different genomes, they have been divided by the estimated genome size.

We classified each metrics using a color code, in order to ease the table reading. Each cell was colored in green if a criteria is fully satisfied (high quality), red if it is not satisfied (low quality), and orange if it is in between (medium quality). We tried to use a criteria that was previously published. The quality criteria, the colour code and the corresponding bibliographic references can be found on Table 3.

**Table 3.** Metrics and thresholds used to evaluate assemblies and assemblers.

|  | low | medium | high | references |
| --- | --- | --- | --- | --- |
| Contig NG50 | < 1 Mb | 1 Mb ≤ 10 Mb | > 10 Mb | (Rhie, McCarthy, et al., 2021) |
| Contig LG50 | > 200 | 50 ≤ 200 | > 200 | |
| k-mer completeness | < 80% | 80% ≤ 95% | > 95% | (Rhie, McCarthy, et al., 2021; Rhie, Walenz, et al., 2020) |
| genome coverage difference | > 40% | 40% ≤ 20% | < 20% | |
| reference assembly coverage difference | > 20% | 20% ≤ 10% | < 10% | |
| BUSCO complete | < 80% | 80% ≤ 95% | > 95% | (Rhie, McCarthy, et al., 2021) |
| BUSCO duplicated | > 5% | 2% ≤ 5% | < 2% | |
| Basepair QV | < 30 | 30 ≤ 50 | > 50 | (Rhie, McCarthy, et al., 2021; Rhie, Walenz, et al., 2020) |
| Assembly errors | > 100 | 20 ≤ 100 | < 100 | |
| memory | > 125GB/Gbp | 32GB/Gbp ≤ 125GB/Gbp | < 32GB/Gbp | |
| time | > 1 week on 8 cores per Gbp | 1 day on 8 cores per Gbp ≤ 1 week on 8 cores per Gbp | < 1 day on 8 cores per Gbp | |

### Read sets

The reads generated by the sequencing protocols have also specifics characteristics presented in Table 4.

**Table 4.** Metrics on long read datasets used in the benchmark.

| Species | technology | # reads | # base pairs | mean length (bp) | coverage |
| --- | --- | --- | --- | --- | --- |
| *Bos taurus* | HiFi | 7,430,931 | 117,560,145,736 | 15,820.4 | 33.30 |
| *Bos taurus* | ONT | 10,144,221 | 156,488,832,730 | 15,426.4 | 44.33 |
| *Bos taurus* | CLR | 14,765,644 | 279,331,710,507 | 18,917.7 | 102.85 |
| *Zea mays* | HiFi | 5,976,900 | 75,601,454,246 | 12,648.9 | 29.42 |
| *Zea mays* | ONT | 12,698,704 | 145,838,781,573 | 11,484.5 | 56.75 |
| *Zea mays* | CLR | 6,114,831 | 130,950,539,094 | 21,415.2 | 50.95 |
| *Coturnix japonica* | HiFi | 1,851,480 | 30,108,307,131 | 16,261.8 | 22.81 |
| *Coturnix japonica* | ONT | 4,147,888 | 65,000,070,596 | 15,670.6 | 49.24 |

The *Bos taurus* datasets have been first published by Eché et al., 2023, whereas the other datasets will be published in other data papers. All datasets are available at EMBL-EBI ENA archive. The corresponding accessions can be found in in Table 5.

**Table 5.** Public accession number of long reads used in the benchmarks.

| Species | technology | accessions |
| --- | --- | --- |
| *Bos taurus* | HiFi | ERR10378054 to ERR10378058 |
| *Bos taurus* | ONT | ERR10386215 to ERR10386219 |
| *Bos taurus* | CLR | ERR10378053 |
| *Zea mays* | HiFi | ERR14085295, ERR14085303, ERR14085304 |
| *Zea mays* | ONT | ERR14056479 to ERR14056485 |
| *Zea mays* | CLR | ERR14085305, ERR14085306 |
| *Coturnix japonica* | HiFi | ERR11528759, ERR11528760 |
| *Coturnix japonica* | ONT | ERR11591487, ERR11591488 |

### Overview of long read de novo assembly methods

Choosing an assembler is not easy because novel software packages are coming out regularly and existing assemblers change version sometimes frequently. The criteria used to chose the assemblers were mainly their actual usage in the community and their efficacy. Canu (Koren et al., 2017) and necat (Y Chen, Nie, et al., 2021) were left out because of the number of computing hours need to assemble a large genome.

Initial long read genome assembly software packages list is given in Table 6.

**Table 6.** Assembler versions and technologies used in the benchmark.

| Assembler | Version | ONT | CLR | HiFi | Reference | Comments |
| --- | --- | --- | --- | --- | --- | --- |
| canu | 2.2 | X | X | X | (Koren et al., 2017) |  |
| flye | 2.9 | ✓ | ✓ | ✓ | (Kolmogorov et al., 2019) | contig polishing |
| hifiasm | 0.16.1 | X | X | ✓ | (Cheng et al., 2022) | read correction |
| ipa | 1.0.3-0 | X | X | ✓ | (Sovic et al., 2022) |  |
| lja | V0.1 | X | X | ✓ | (Bankevich et al., 2022) |  |
| necat | 20200803 | X | X | X | (Y Chen, Nie, et al., 2021) |  |
| NextDenovo | 2.5.0 | ✓ | ✓ | X | (Hu et al., 2023) |  |
| raven | 1.8.1 | ✓ | X | X | (Vaser and Šikić, 2021) |  |
| shasta | 0.5.1 | ✓ | ✓ | X | (Shafin et al., 2020) |  |
| wtdbg2 | 2.5 | ✓ | ✓ | X | (Ruan and Li, 2019) |  |

Not all data types, species, assembler combinations could be run because CLR data were not available for *Coturnix japonica*. Some assemblers are data specific: for example, hifiasm, ipa, lja cannot assemble ONT nor CLR reads. Finally, some combinations did not terminate correctly, despite several tries: shasta on *Zea mays* ONT reads, and ipa on HiFi reads for the same species.

**Table 7.** Metrics for *Bos taurus*. Description of abbreviations: NG50: NG50 in MB, comp: completeness, cov.: genome coverage, B. C: BUSCO C, B. D: BUSCO D, err.: number of errors, mem.: memory in GB.

| type | assembler | NG50 | LG50 | comp. | rel. size | cov. | B. C | B. D | QV | err. | time | mem. |
| --- | --- | --- | --- | --- | --- | --- | --- | --- | --- | --- | --- | --- |
| Ref. |  | 82.40 | 16 | NA | -26% | NA | 96% | 2% | NA | NA | NA | NA |
| ONT | flye | 26.38 | 33 | 93% | -25% | 99% | 86% | 1% | 31 | 25 | 33d | 88.3 |
| ONT | NextDenovo | 58.92 | 21 | 92% | -25% | 99% | 85% | 2% | 31 | 37 | 108d | 16.9 |
| ONT | raven | 12.92 | 70 | 37% | -26% | 7% | 9% | 0% | 15 | 14893 | 6d | 27.5 |
| ONT | shasta | 18.13 | 55 | 91% | -26% | 98% | 81% | 1% | 29 | 779 | <1d | 205.3 |
| ONT | wtdbg2 | 14.64 | 56 | 87% | -23% | 97% | 75% | 1% | 25 | 146 | 62d | 97.3 |
| CLR | flye | 12.29 | 66 | 96% | -25% | 99% | 95% | 2% | 36 | 871 | 55d | 90.3 |
| CLR | NextDenovo | 42.98 | 23 | 95% | -25% | 99% | 92% | 2% | 34 | 571 | 358d | 16.4 |
| CLR | raven | 10.69 | 76 | 14% | -22% | 0% | 1% | 0% | 12 | 112914 | 3d | 29.8 |
| CLR | shasta | 0.09 | 6843 | 54% | -43% | 16% | 35% | 0% | 19 | 213 | <1d | 127.2 |
| CLR | wtdbg2 | 9.82 | 80 | 93% | -25% | 97% | 90% | 1% | 29 | 393 | 25d | 71.6 |
| Hifi | flye | 0.72 | 998 | 98% | -2% | 99% | 94% | 6% | 49 | 58 | 20d | 62.8 |
| Hifi | hifiasm | 76.68 | 17 | 96% | -9% | 100% | 96% | 2% | 66 | 13 | 12d | 47.3 |
| Hifi | ipa | 32.46 | 30 | 96% | -16% | 99% | 96% | 3% | 56 | 67 | 15d | 9.7 |
| Hifi | lja | 1.09 | 942 | 100% | 62% | 100% | 91% | 26% | 50 | 2 | 18d | 77.2 |
| Hifi | raven | 2.80 | 301 | 92% | -24% | 99% | 87% | 1% | 45 | 342 | 1d | 23.1 |
| Hifi | wtdbg2 | 18.30 | 47 | 93% | -27% | 96% | 94% | 2% | 41 | 52 | 11d | 43 |

**Table 8.**
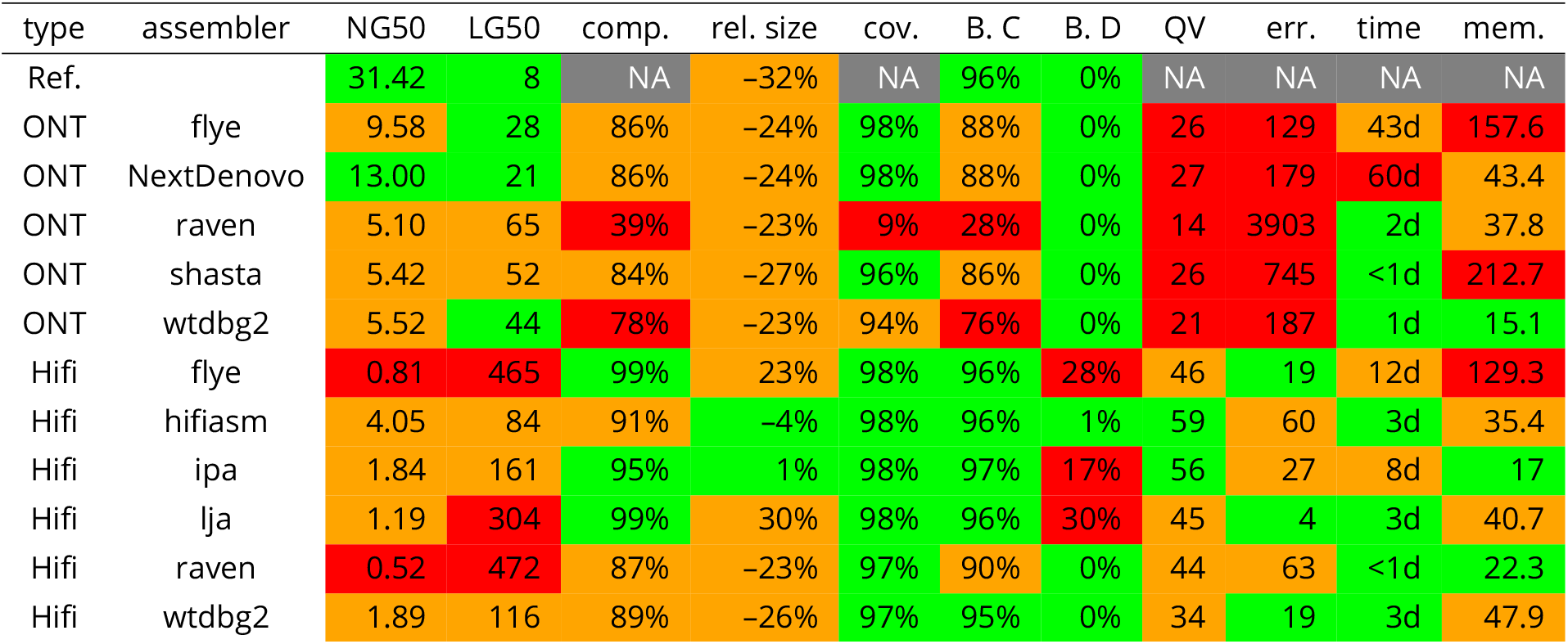
Metrics for *Coturnix japonica*.

| type | assembler | NG50 | LG50 | comp. | rel. size | cov. | B. C | B. D | QV | err. | time | mem. |
| --- | --- | --- | --- | --- | --- | --- | --- | --- | --- | --- | --- | --- |
| Ref. |  | 31.42 | 8 | NA | -32% | NA | 96% | 0% | NA | NA | NA | NA |
| ONT | flye | 9.58 | 28 | 86% | -24% | 98% | 88% | 0% | 26 | 129 | 43d | 157.6 |
| ONT | NextDenovo | 13.00 | 21 | 86% | -24% | 98% | 88% | 0% | 27 | 179 | 60d | 43.4 |
| ONT | raven | 5.10 | 65 | 39% | -23% | 9% | 28% | 0% | 14 | 3903 | 2d | 37.8 |
| ONT | shasta | 5.42 | 52 | 84% | -27% | 96% | 86% | 0% | 26 | 745 | <1d | 212.7 |
| ONT | wtdbg2 | 5.52 | 44 | 78% | -23% | 94% | 76% | 0% | 21 | 187 | 1d | 15.1 |
| Hifi | flye | 0.81 | 465 | 99% | 23% | 98% | 96% | 28% | 46 | 19 | 12d | 129.3 |
| Hifi | hifiasm | 4.05 | 84 | 91% | -4% | 98% | 96% | 1% | 59 | 60 | 3d | 35.4 |
| Hifi | ipa | 1.84 | 161 | 95% | 1% | 98% | 97% | 17% | 56 | 27 | 8d | 17 |
| Hifi | lja | 1.19 | 304 | 99% | 30% | 98% | 96% | 30% | 45 | 4 | 3d | 40.7 |
| Hifi | raven | 0.52 | 472 | 87% | -23% | 97% | 90% | 0% | 44 | 63 | <1d | 22.3 |
| Hifi | wtdbg2 | 1.89 | 116 | 89% | -26% | 97% | 95% | 0% | 34 | 19 | 3d | 47.9 |

**Table 9.** Metrics for *Zea mays*.

| type | assembler | NG50 | LG50 | comp. | rel. size | cov. | B. C | B. D | QV | err. | time | mem. |
| --- | --- | --- | --- | --- | --- | --- | --- | --- | --- | --- | --- | --- |
| Ref. |  | 217.87 | 5 | NA | -20% | NA | 95% | 14% | NA | NA | NA | NA |
| ONT | flye | 1.67 | 379 | 78% | -16% | 49% | 94% | 13% | 20 | 72 | 111d | 135.7 |
| ONT | NextDenovo | 31.31 | 23 | 77% | -19% | 39% | 94% | 12% | 20 | 330 | 244d | 25.7 |
| ONT | raven | 0.11 | 6418 | 23% | -30% | 0% | 11% | 0% | 12 | 0 | 8d | 39.2 |
| ONT | wtdbg2 | 0.12 | 6219 | 65% | 37% | 22% | 84% | 12% | 15 | 0 | 17d | 156.1 |
| CLR | flye | 4.12 | 158 | 98% | -18% | 60% | 98% | 15% | 42 | 138 | 145d | 133.8 |
| CLR | raven | 0.75 | 1000 | 24% | -5% | 0% | 4% | 0% | 12 | 44982 | 6d | 31.2 |
| CLR | shasta | 0.22 | 2647 | 87% | -31% | 47% | 86% | 9% | 23 | 429 | <1d | 189.6 |
| CLR | wtdbg2 | 0.17 | 4617 | 91% | 89% | 50% | 88% | 20% | 17 | 2 | 13d | 126.7 |
| Hifi | flye | 10.39 | 63 | 98% | -17% | 55% | 98% | 15% | 63 | 13 | 16d | 200.3 |
| Hifi | hifiasm | 53.62 | 13 | 98% | -15% | 54% | 98% | 15% | 67 | 5 | 7d | 46 |
| Hifi | lja | 26.53 | 28 | 98% | -17% | 60% | 98% | 15% | 58 | 1 | 38d | 63.3 |
| Hifi | raven | 0.00 | 21551 | 57% | -71% | 25% | 75% | 6% | 42 | 0 | 2d | 24 |
| Hifi | wtdbg2 | 0.07 | 8097 | 92% | -29% | 54% | 96% | 15% | 36 | 0 | 59d | 103.2 |

### Experimental setup

The benchmark was performed on 2.1GHz Intel Xeon E5-2683 v4 CPUs, on CentOS 7 Linux distributions.

## Results

The names of the assemblies presented in this section will join the sequencing protocole (HiFi, CLR, ONT) and the assembler (flye, hifiasm, ipa, lja, NextDenovo, raven, shasta, or wtdbg2).

### *Bos taurus* results

We can note at first glance that three assemblies —namely ONT raven, CLR raven, and CLR shasta— are not satisfactory, observing several metrics such as completeness, genome coverage, BUSCO C, QV, and number of errors. These assemblies are significantly too short to be useful, and will not be discussed in the rest of the analysis.

NG50 and LG50, together show that HiFi lja or HiFi raven provide the most fragmented assemblies. On the other end, HiFi hifiasm, HiFi ipa, CLR NextDenovo, ONT NextDenovo and ONT flye output the least fragmented genome assemblies.

A striking result is that the genome coverage is very high for all remaining assemblies, meaning that almost all the reference genome is covered by the remaining assemblies. However, completeness, which is based on k-mers instead of alignments, gives another picture: the completeness of most ONT and CLR assemblies (with the notable exception of CLR flye), as well as HiFi raven and HiFi wtdbg2, is below 95%. This suggests that ONT and CLR assemblies, as expected, contain small errors, although flye corrects these errors in its pipe-line. This result is confirmed by the BUSCO C score, the QV, and the number of errors: best scores are given to HiFi assemblies, while CLR assemblies have slightly better scores than ONT. Of note, the error metrics is sensitive to any kind of mutation, even point mutations, which could give rise to frame-shift or early stop codons in the translated proteins.

This is also confirmed, to some extent, by the number of errors, which is higher for CLR and ONT data, compared to HiFi.

Another interesting result is the relative size, compared to cytology analysis. Most assemblies, including the reference one, are significantly shorter than the predicted size. This could suggest that the repeated fraction of the genome is poorly assembled. HiFi flye, HiFi hifiasm and to a lesser degree, HiFi ipa are the only assemblies which sizes are close to the goal. However, the assembly size of HiFi flye is somewhat misleading because it has a BUSCO D of 6% clearly suggesting that it is partly duplicated. If we link this result with the low NG50, and the high LG50, this suggests that HiFi flye kept haplotypes apart, cutting the assembly each time two alleles where present and keeping both of them.

It is also fruitful to compare the results produced by the same tool, and different data. NextDenovo, for instance, has been used on ONT and CLR data. NextDenovo ONT gives a more contiguous assembly (NG50: 59M vs 43M, LG50: 21 vs 23). However, NextDenovo CLR shows less errors (BUSCO C: 92% vs 85%, QV: 34 vs 31), and reflects the reads better (completeness: 95% vs 92%). Other tools where used with the three technologies: flye, raven and wtdbg2. They also confirm that ONT assemblies are longer, but more error-prone than CLR assemblies. There are fewer errors in HiFi assemblies, when compared to CLR assemblies (for HiFi wtdgb2 and CLR wtdbg2, BUSCO C: 94% vs 90%, QV: 41 vs 29). The assembly size however, is not directly linked to the type of data: HiFi wtdbg2 assembly has a longer NG50 (18M vs 10M) and a smaller LG50 (47 vs 80), but we observe the opposite for HiFi raven (NG50: 3M vs 13M, LG50: 301 vs 70) and HiFi flye (NG50: 700K vs 26M, LG50: 998 vs 33). This is a probable consequence of the fact that flye and raven separate haplotypes.

### *Coturnix japonica* results

Assemblies of *Coturnix japonica* are much more fragmented than *Bos taurus*, and the NG50 ranges between 500k (for HiFi raven and flye) to 13M (for CLR NextDenovo). The NG50 is actually sensible to the genome size, and the size of the *Coturnix japonica* genome is much smaller than the size of the *Bos taurus* genome. Quail has also few large chromosomes and a large number of micro-chromosomes which limits the possible NG50 size.

Akin to the previous data-set, BUSCO C score, QV, and the number of errors, show that ONT assemblies contain more errors, compared to HiFi. Moreover, assembly metrics are longer with ONT data (longest NG50 is 10M for ONT flye, and shortest is 500k for HiFi raven).

The relative size shows that most tools only assemble part of the genome, with score around *−*24%, which is still higher than the reference assembly (*−*32%). HiFi flye, HiFi hifiasm, HiFi ipa, and HiFi lja are the only exceptions. Flye lja and, to a lesser degree, HiFi ipa have positive relative sizes with BUSCO D scores up to 30%. This suggests that both haplotypes have been included in the assemblies. HiFi hifiasm is the only assembly with a good relative size, and a modest BUSCO D score of 1%. Its completeness of 91%, which is lower than other assemblies (e.g. HiFi flye, HiFi ipa and HiFi lja) may confirm that only one allele was kept in the final assembly.

### *Zea mays* results

Several assemblers including ONT raven, ONT wtdbg2, CLR raven, HiFi raven, failed on this dataset. They produce very short assemblies. The genome coverage of raven is really low (down to 0%) for both ONT and CLR read types. wtdbg2 NG50 did not exceed 170K on all read types. ONT NextDenovo, as well as HiFi flye, hifiasm, and lja, provided rather contiguous assemblies, with NG50 greater than 10M.

A first observation is that the genome coverage never exceeds 60%. At a first glance, this could imply that roughly half of the reference genomes is not syntenic with the assembled genome. However, the reason could be elsewhere. In order to compute the genome coverage, the assembled genome is mapped to the reference genome. Two copies of the same repeat in the assembled genome will like map to the same copy in the reference. As a consequence, the unmapped copy of the reference genome will be detected as uncovered, and reported as such.

The BUSCO D is also very low. However, this is most probably due to a whole genome duplication which only happened in this species (Schnable et al., 2009). As a consequence, the private copied genes are flagged as duplicated by BUSCO.

Above all, shasta assemblies seem problematic, since the CLR and ONT assemblies do not resemble the reference assembly (genome coverage of 0%). Confirming this observation, the relative size is almost always negative, except for ONT and CLR wtdbg2. On the other side, CLR flye, as well as HiFi flye, hifiasm, and lja, show a completeness of 98%, indicating that almost all the reads have been used in the assembly. This may suggest that the reference genome is very divergent from the assembled one, or that repeats are poorly assembled. Another striking result is that, although the assemblies are smaller than expected, the BUSCO C score can reach 98% (for CLR flye, as well as HiFi flye, hifiasm, and lja), which is higher than the BUSCO C score for the reference genome of 95%. This suggests that essential genes are found in these assemblies.

The BUSCO D score, however, is not less than 9%, including the reference genome except for raven assemblies. This strongly suggests that the genome is highly repeated, and that repetitions include genes.

As previously observed, HiFi assemblies include less errors than other assemblies.

## Discussion

### Assembly accuracy

#### Choice of the sequencing strategy

The type of sequenced reads is critical to obtain a high quality assembly: HiFi reads deliver better assemblies, with longer NG50, smaller LG50, reflecting reference genomes and reads better, with better BUSCO scores. CLR data, on our benchmarks, seem to be slightly better than ONT reads, especially when used with flye, which corrects read errors.

The assemblers also provide very distinct results, depending on the sequencing strategy. Hifiasm, only applicable to HiFi data, seems to provide the best results. Flye provides the best results for CLR reads, and NextDenovo, the best results for ONT reads.

#### Impact of the genome

Complexity of sequenceed genomes does also impact assembly quality. Assemblies appear to be more fragmented when genomes are heterozygous. As shown in the *Bos taurus* example, some tools may cut the assembly each time they observe several alleles, and possibly report them. This may obviously over-estimate the size of the assembly, especially if alternative allele are not appropriately handled. New pipe-lines, which include verkko (Rautiainen et al., 2023), are specifically designed to assemble diploid genomes, and provide haplotyped assemblies.

Genomes with many repetitions may also be poorly assembled. The different copies of the repetitions may be “collapsed” into a single sequence. In order to solve this problem, assemblers relies on small differences that appear between copies. Discriminating between real differences and sequencing errors is a key feature in order to correctly assemble reads.

For this, the tool should be able to differentiate real differences, and sequencing errors. This is especially difficult for ONT reads, or uncorrected CLR reads, which accumulate errors. For long, highly repetitive, regions, consortium for telomere-to-telomere assemblies (Nurk et al., 2022) suggests the use of ultra-long reads, with size larger than 100kb, in order to assemble them.

#### Time and memory usage

Concerning time, NextDenovo is by far the slowest tool, and flye usually is the second slowest. shasta is the fastest tool, and raven the second fastest. ipa and lja usually are between the two previous groups. wtdbg2 shows an irregular profile: it can be almost as fast as raven (*Zea mays* with ONT data), and slower than flye (*Bos taurus* with ONT data). This tool seems to be highly dependant on both the species and the data. flye also seems to be much faster on HiFi data.

Concerning memory, shasta requires more memory than other tools, flye is usually second, and wtdbg2, usually third. NextDenovo, raven, and ipa require less memory than other tools.

NextDenovo and shasta exhibit an opposite profile: the former requires more time and less memory, the latter requires less time and more memory. This trade-off is most likely the consequence of implementation choices.

Raven and ipa are both fast, and require less memory than other tools. On the opposite, flye is rather slow, and requires more memory than most other tools.

#### Metrics

Defining a high-quality assembly is straight-forward: it accurately provides the series of nucleotides that form each chromosome. However, there are many types of low-quality assemblies: missing regions, duplications, etc. The metrics we used make it possible to assess the different possible types of errors in the assemblies. Obviously, most metrics evaluate several criteria at the same time: the NG50 will be low if the assembly is fragmented, and also if the genome is incomplete. The BUSCO C, for instance, may be used to evaluate unused read sequences, duplication of sequences, and short sequencing errors.

Some metrics should be considered with extra caution, especially when the genome is highly repeated. The *Zea mays* genome, for instance, shows very poor genome coverage and BUSCO D results. A telomere-to-telomere assembly (J Chen et al., 2023), which has been published after this analysis was carried out, informed that the genome contains many tandem (near exact) repeats, highly repeated centromeric, telomeric and subtelomeric regions, and numerous satellite and super-long simple-sequence-repeat arrays. This, together with a recent whole genome duplication, are crucial information that make it possible to correctly interpret poor metrics.

## Acknowledgements

We are grateful to the GenoToul bioinformatics platform Toulouse Occitanie (Bioinfo Genotoul, https://doi.org/10.15454/1.5572369328961167E12) for providing help and/or computing and/or storage resources.

We thank members of SeqOccIn Consortium for their feedback before and during this study.

## Data, script, code, and supplementary information availability

Assemblies are available on a dedicated repository https://doi.org/10.57745/63UXPG. Tool versions, and commands used are available on https://forgemia.inra.fr/seqoccin/assembler-benchmarking. Public accessions of the long read data sets used for this study are presented in Table 5.

